# Discovery of a novel inhibitor of macropinocytosis with antiviral activity

**DOI:** 10.1101/2023.10.25.563967

**Authors:** Bartlomiej Porebski, Wanda Christ, Alba Corman, Martin Haraldsson, Myriam Barz, Louise Lidemalm, Maria Häggblad, Juliana Ilmain, Shane C. Wright, Matilde Murga, Jan Shlegel, Erdinc Sezgin, Gira Bhabha, Volker M Lauschke, Miguel Lafarga, Jonas Klingström, Daniela Huhn, Oscar Fernandez-Capetillo

**Author notes:** **Lead Contact:** Oscar Fernandez-Capetillo, Science for Life Laboratory, Division of Genome Biology, Department of Medical Biochemistry and Biophysics, Karolinska Institute, S-171 21 Stockholm, Sweden, Twitter: @KP_Twitt_llo.

## Abstract

Several viruses hijack various forms of endocytosis in order to infect host cells. Here, we report the discovery of a new molecule with antiviral properties that we named virapinib, which limits viral entry by macropinocytosis. The identification of virapinib derives from a chemical screen using High-Throughput Microscopy, where we identified new chemical entities capable of preventing infection with a pseudotype virus expressing the spike (S) protein from SARS-CoV-2. Subsequent experiments confirmed the capacity of virapinib to inhibit infection by SARS-CoV-2, as well as by additional viruses, such as Monkeypox virus and TBEV. Mechanistic analyses revealed that the compound inhibited macropinocytosis, limiting this entry route for the viruses. Importantly, virapinib has no significant toxicity to host cells. In summary, we present a new molecule that inhibits viral entry via the endocytic route, offering a new alternative to prevent viral infection.

## INTRODUCTION

Since its onset in 2020, the Covid-19 pandemic has caused close to 7 million deaths across all continents and has led to major disturbances in the world economy (https://covid19.who.int/). The disease is caused by Severe Acute Respiratory Syndrome Coronavirus 2 (SARS-CoV-2), which belongs to the *betacoronavirus* genus of the *coronavirus* family of positive-sense single-stranded RNA viruses. Several human coronaviruses are endemic and normally cause mild respiratory infections. However a few such as MERS-CoV (Middle east respiratory syndrome virus), SARS-CoV and SARS-CoV-2 can cause an acute respiratory syndrome with severe lung injuries and inflammatory responses^1,2^. The SARS-CoV-2 surface is covered with the eponymous ‘crown’ formed by the Spike (S) glycoprotein, which binds to the host-cell receptor, Angiotensin Converting Enzyme 2 (ACE2)^3^.

Due to its profound impact, there has been a global effort to develop therapies to protect against Covid-19, which to a large extent was made possible by RNA-based vaccines. In addition, counteracting the cytokine storm with the steroid dexamethasone has proven a successful strategy to reduce Covid-19-related deaths^4^. Regarding the development of antivirals that limit the infection step or that target SARS-CoV-2, to date, only two drugs targeting the virus have received medical approval: paxlovid (targeting the viral protease) and remdesivir (targeting the RNA-dependent RNA polymerase)^5^. Although numerous drug-repurposing approaches have been reported, none have reached clinical practice^6^.

An alternative to target the virus is to target its host cells, which is argued to be less susceptible to resistance mechanisms and potentially serve as a strategy to prevent infection by multiple viruses^7^. Successful examples of this approach include CCR5 antagonists for HIV and interferon for hepatitis B or C (HCV and HBV) viruses. To date, no host-targeting antiviral agents have been approved against SARS-CoV-2. By conducting a phenotypic chemical screen using pseudotype viruses expressing the S protein of SARS-CoV-2, we here report the identification of a new chemical entity that limits viral entry by endocytic pathways for several viruses including SARS-CoV-2.

## RESULTS

### A pseudovirus-based screening assay for SARS-CoV-2 infection

To identify small molecules that inhibit SARS-CoV-2 entry, we used lentiviral pseudotyped viruses that expressed a cytomegalovirus (CMV)-driven Enhanced Green Fluorescent Protein (EGFP) as well as a Flag-tagged S protein from SARS-CoV-2 (**Fig. 1A**). The S protein lacked the last 19 aminoacids, which was shown to facilitate membrane incorporation by deleting an endoplasmic reticulum (ER)-retention signal^8^. S-enveloped pseudoviral particles (PV^S^) were produced in HEK 293T cells, as confirmed by western blot (WB) analysis of supernatants **(Fig. 1B)**. To facilitate infection by PV^s^, we generated a clone of HEK 293T cells expressing the ACE2 receptor (293T^ACE2^) **(Fig. 1C)**. Quantification of infection rates by high-throughput microscopy indicated that 1.91% of 293T^ACE2^ cells were successfully infected with the PV^S^ **(Fig. 1D, E)**. Furthermore, the number of infected cells was reduced approximately 8-fold by treatment with E-64d, a Cathepsin inhibitor that inhibits infection by SARS-CoV-2 and S-enveloped pseudoviruses^3,8^. Subsequent repeats of the experiment confirmed its reproducibility, and the assay window was sufficient to conduct chemical screening.

**Figure 1.**
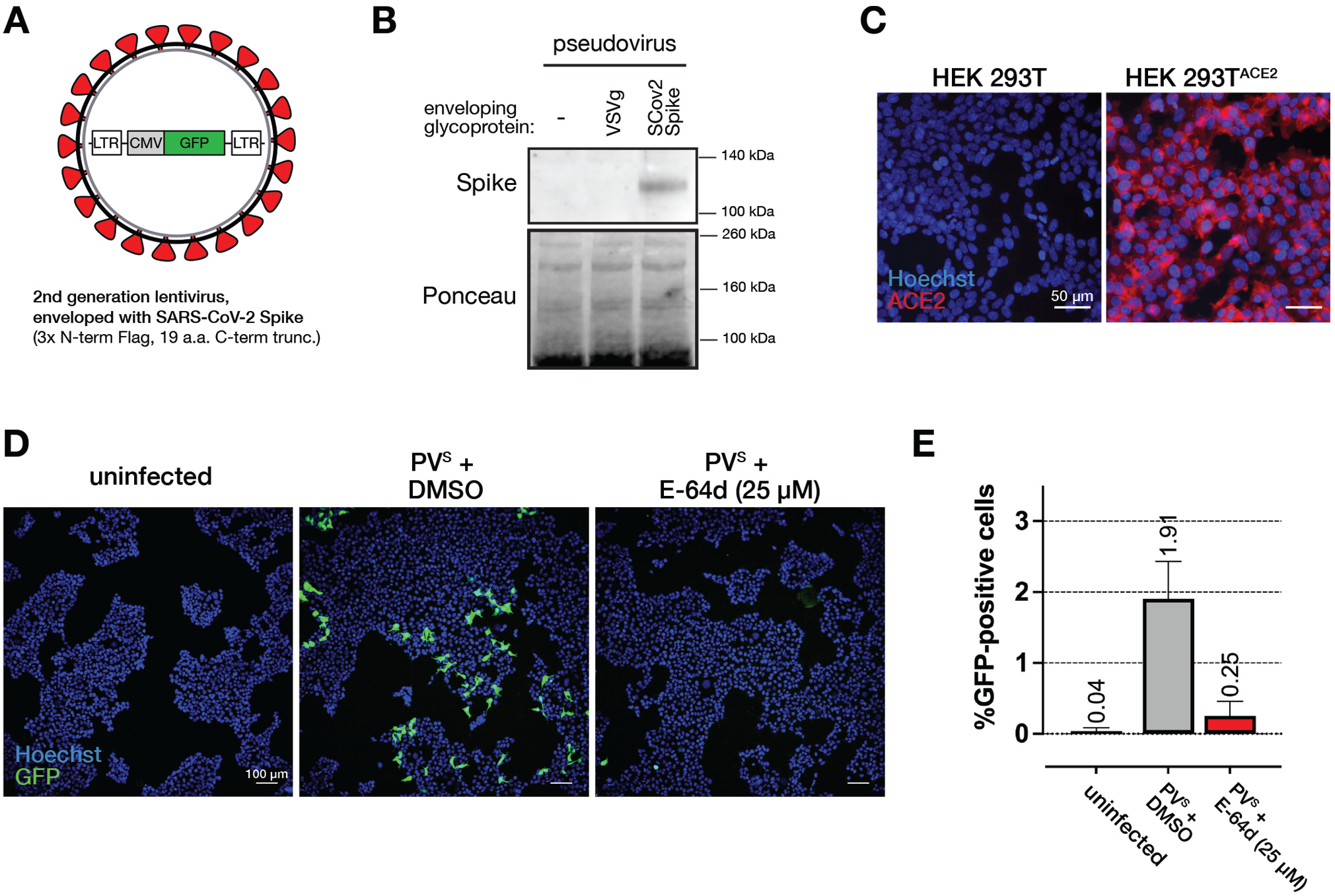
A S-pseudotyped lentivirus to model SARS-CoV-2 infection. (**A**) Schematic of the S-pseudotyped virus (PV^S^) expressing GFP. (**B**) WB analysis of viral particle-rich supernatants. The image of the Ponceau gel is shown to illustrate equivalent loading controls. (**C**) Representative images of wild-type (left) or HEK 293T^ACE2^ cells (right) stained with Hoechst 33342 (blue), to visualize nuclei, and anti-ACE2 antibody (red). Scale bar (white) represents 50 μm. (**D**) Representative images of the screening assay. HEK 293T^ACE2^ cells either uninfected (left), infected with PV^S^ and treated with DMSO (middle) or infected with PV^S^ and treated with Cathepsin inhibitor E-64d (right). Infection was detected on the basis of GFP (green) expression. Hoechst 33342 (blue) was used to visualize nuclei. Cells were treated for 6 h, followed by the addition of PV^S^ and microscopic analysis 72 h after the infection. Scale bar (white) represents 100 μm. (**E**) Quantification of the experiment represented in (**D**). For each condition, 3 separate wells were imaged, each containing ∼4000 cells.

### Identification of small molecules that inhibit infection by PV^S^

Having established the assay, we next performed a chemical screen to identify novel chemical entities that limit infection by PV^S^. As a screening library, we used a set of 1,008 molecules predicted to have properties that could impair protein-protein interactions (see Methods). The choice was made on the basis that SARS-CoV-2 entry is facilitated by protein-protein interactions and to discover new chemistry beyond molecules that are already being investigated as antivirals. In the screening pipeline **(Fig. 2A)**, 293T^ACE2^ cells seeded in 384-well plates were first treated with the compounds from the chemical library (25 μM) for 6 h, followed by the addition of PV^S^ and a subsequent 72 h incubation to allow for EGFP expression. After fixation, DNA was stained with Hoechst 33342 and cells imaged using Hight-Throughput Microscopy (HTM). Infection rates were calculated as the ratio of EGFP-positive cells to the total number of nuclei per well, and all data were normalized to the infection rates observed in wells treated with DMSO (negative control). The raw infection rate across all plates was 3.11% **(Fig. S1A)**. Treatment with E-64d led to an approximately 10-fold decrease in the infection rate **(Fig. 2B).** For hit calling, we selected two criteria: hit compounds had to decrease the infection rate below 20% and show minimal toxicity to host cells. Three compounds met these criteria, and their effects were confirmed by individual inspection of the images of each well **(Fig 2C, D)**.

**Figure 2.**
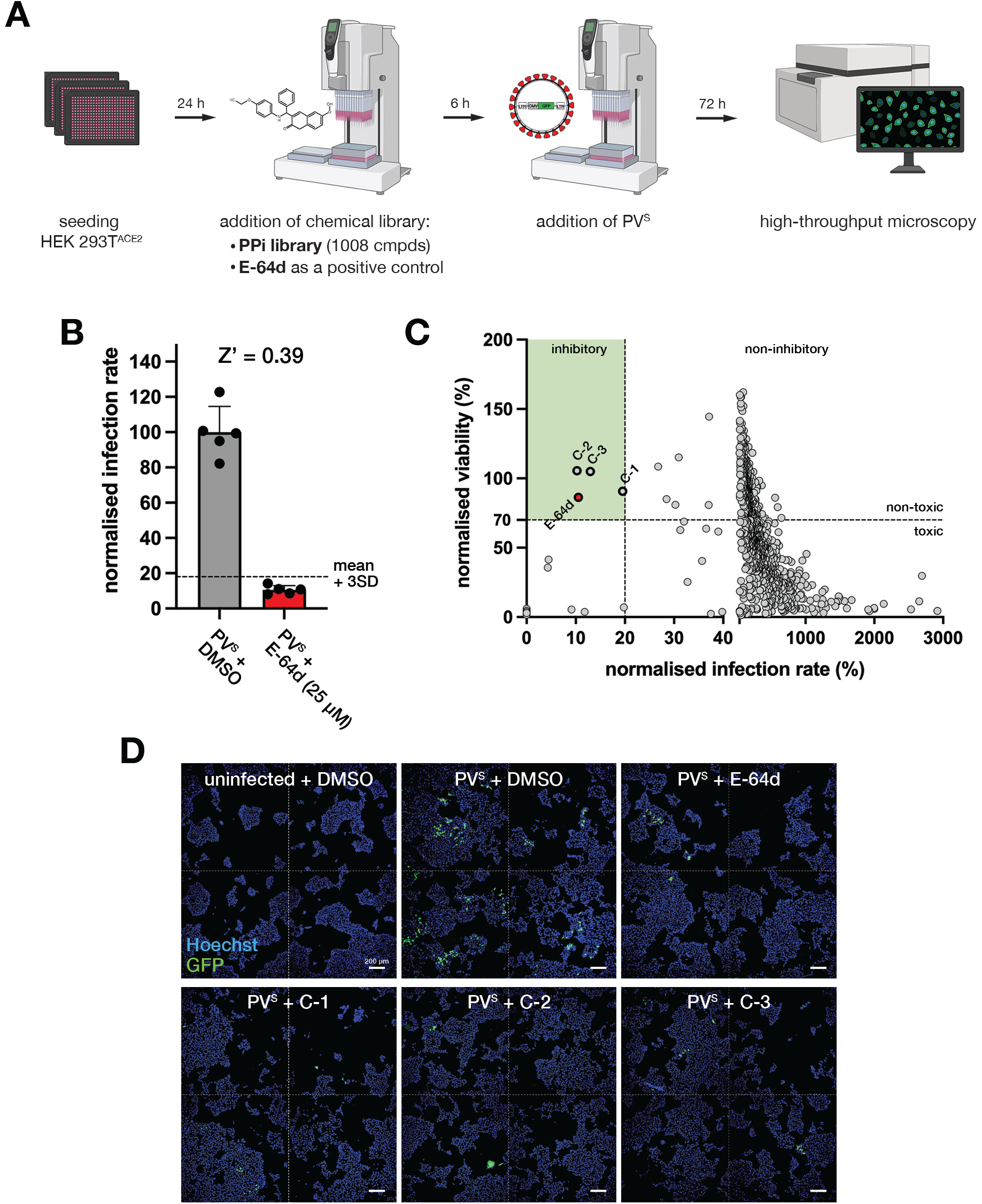
A chemical screen for inhibitors of PV^s^ infection. (**A**) Schematic of the screening pipeline. The chemical library (1,008 compounds from a set designed to inhibit protein protein-interactions: PPi) as well as E-64d were added for 6 h before infection. The final concentration of compounds was 25 μM after addition of the virus. (B) Quality control and assay window of the screen. Each data point represents the mean infection rate calculated from 8 wells within each plate. The Z-prime factor was calculated for the entire plate set. The dashed line marks the mean value + 3 standard deviations (SD) from wells treated with E-64d (25 μM). (**C**) Scatter plot representing the viability and infection rate for each compound in the screen. Dashed lines mark threshold values defined for hit calling; namely limited toxicity (>70% viability) and significant inhibition of infection (<20%). (**D**) Representative microscopy images of wells from the screen (uninfected or infected and treated with DMSO, E-64d or each of the three hits). Hoechst 33342 (blue) was used to visualize nuclei. Dashed lines separate different imaging fields from each well. Scale bar (white) represents 200 μm.

To validate our screening data, we re-tested the three compounds in 293T^ACE2^ cells using the same assay in a dose-response experiment. Although all compounds decreased infection rates, two (C-1 and C-2; structures provided in **Fig. 3A**) showed a dose-dependent effect at the tested concentrations (**Fig. 3B**). Notably, none of the compounds prevented infection with a pseudotype virus expressing the envelope glycoprotein (G) from Vesicular Stomatitis Virus (VSVg), thereby discarding a general effect on host cells or on infection with lentiviral particles (**Fig. 3C**). Furthermore, we confirmed that the compounds did not lower the expression of ACE2 in 293T^ACE2^ cells, nor did they affect EGFP expression in an independent cell line (**Fig. S1B, C**). Although all 3 compounds showed interesting properties in these PV models, the chemical properties of C-2 hampered further development of this chemical series and C-3 failed to show antiviral activity in following assays with replicating virus. Therefore, we selected C-1, which we named virapinib, for the rest of the study.

**Figure 3.**
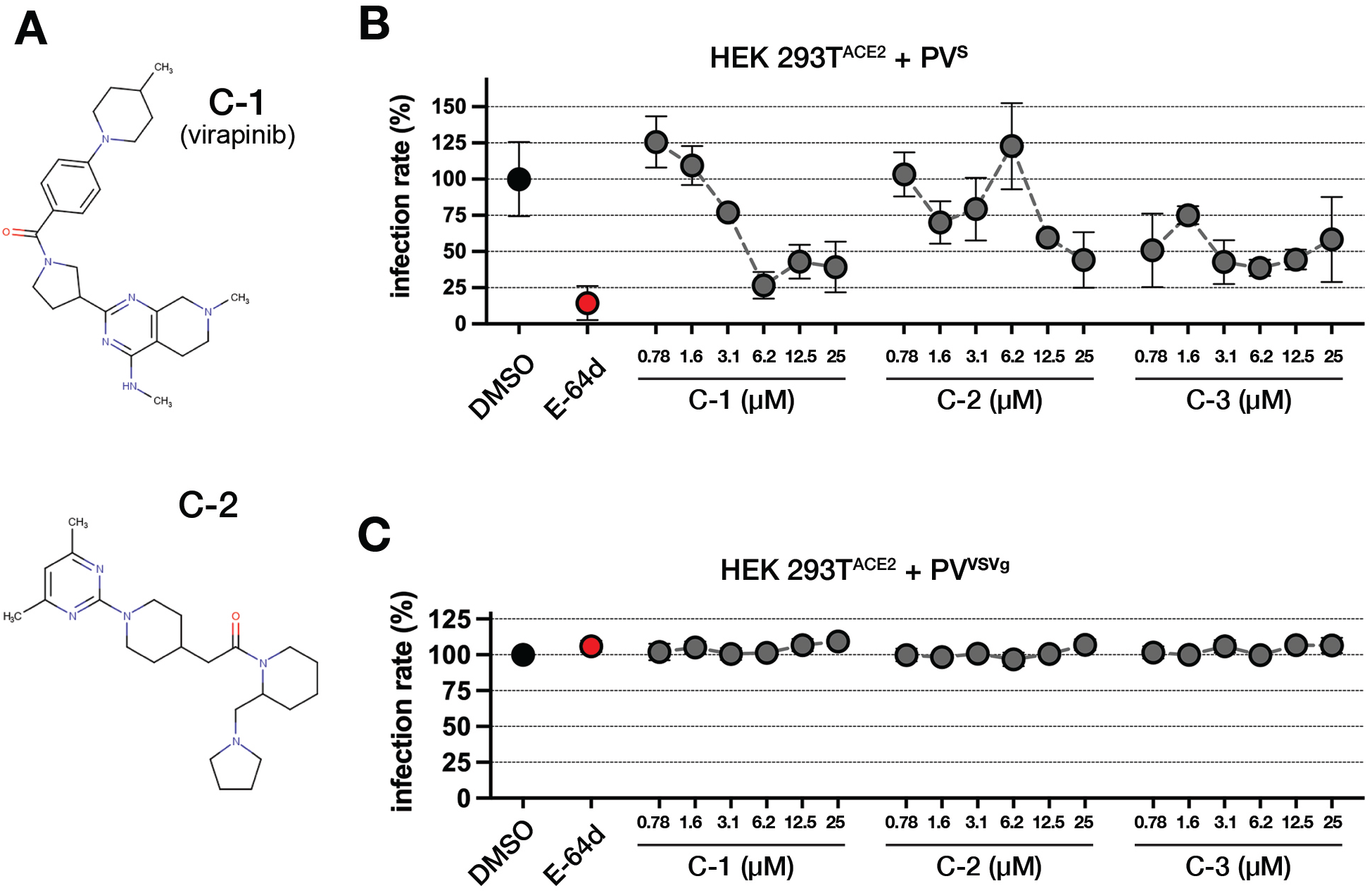
Dose-response analysis of hit compounds in PV^s^ infection. (**A**) Chemical structures of C-1 (virapinib) and C-2. (**B**) Dose response validation of the inhibitory effect of C-1, C-2 and C-3 against PV^s^ infection. Plotted values represent the mean ± SD of 3 wells, each containing ∼4000 cells. E-64d was used as an inhibition control (25 μM). (**B**) Dose response of the effect of C-1, C-2 and C-3 against VSVg-pseudotyped lentiviral infection. Plotted values represent the mean ± SD of 3 wells, each containing ∼4000 cells.

### Virapinib prevents infection with SARS-CoV-2

Next, we tested if virapinib could limit infection with the actual SARS-CoV-2 virus. To focus on the infection step, cells were treated with the compound for 6 h before addition of the virus. After washing off the virus, cells were incubated for 24 h, followed by HTM-dependent quantification of the expression of the N protein from SARS-CoV-2. Using this setup, we first tested the efficacy of virapinib against the ancestral strain of the virus in monkey Vero E6 cells, which are kidney cells from African green monkey origin that have been widely used as host cells for testing viral infections. Consistent with previous studies, Vero E6 cells were efficiently infected with SARS-CoV-2 **(Fig. S2A),** and infection was nearly completely abrogated by E-64d (**Fig. 4A)**^3,8^. In agreement with our results with PV^s^, virapinib presented a dose-dependent effect in limiting SARS-CoV-2 infection in Vero E6 cells and had no detectable toxic or cytostatic effects in this cell line **(Fig. 4A, Fig. S2B)**. Interestingly, while infection rates were reduced by the compound, infection was limited to a few cell clusters where the levels of N-protein were higher than those in control infections, an observation that is also seen with E-64d **(Fig. S2C)**.

**Figure 4.**
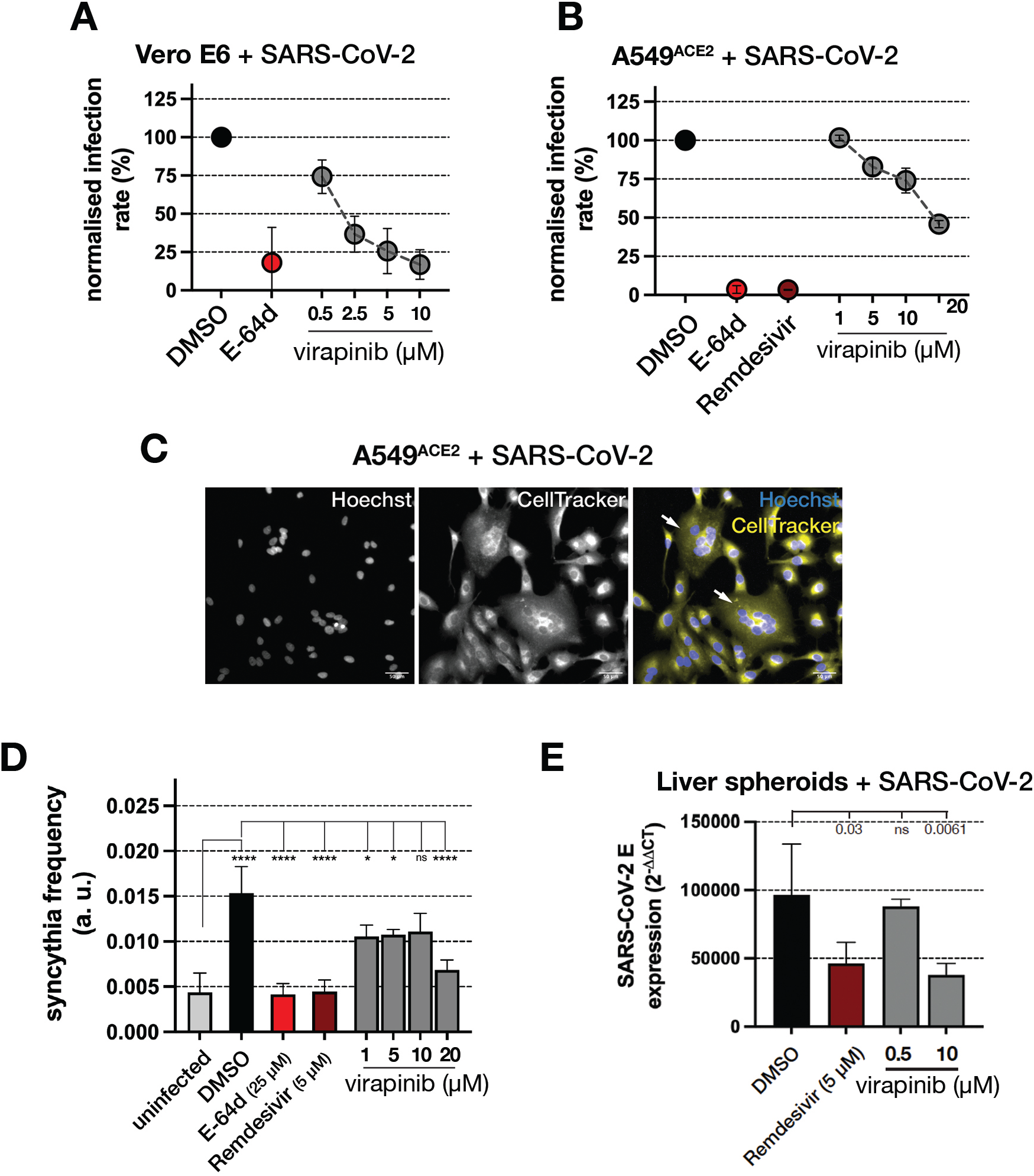
Virapinib inhibits infection with replication-competent SARS-CoV-2. (**A**) Dose-response of virapinib against ancestral SARS-CoV-2 infection in Vero E6 cells. Cells were pre-treated with the compound for 6 h, followed by 1h infection. Cells were fixed at 24hpi and infection rates were calculated by HTM quantifying cells expressing the N-protein from SARS-CoV-2. Each data point represents the mean ± S.D. of 3 wells, each containing ∼15000 cells. E-64d was used as an inhibition control (25 μM). (**B**) As in (A), but with the SARS-CoV-2 infection performed in A549^ACE2^ cells. E-64d (25 μM) and remdesivir (5μM) were used as inhibition controls. (**C**) Representative image of SARS-CoV-2-infected A549^ACE2^ cells illustrating the formation of multi-nucleated syncytia (white arrows). The cell perimeter was labelled using Cell Tracker (yellow). Hoechst 33342 (blue) was used to visualize nuclei. (**D**) Quantification of syncytia formation. Syncytia were defined as cells with continuous cytoplasm and containing more than 2 nuclei, and syncytia frequency was defined as number of syncytia divided by the total number of cells. Plotted data represent mean syncytia frequency from 3 independent samples. ****, *p* < 0.0001; *, *p* < 0.05; ns, non-significant. (**E**) Representative data from the effect of DMSO, remdesivir or virapinib in SARS-CoV-2 infection of liver spheroids. Viral loads where quantified by RT-PCR analysis of SARS-CoV-2 E protein expression. Data represents mean ± S.D. of 3 replicate reactions. Statistical analysis was done with a t-test. Ns, non-significant.

We then aimed to validate these findings in a human cell line. To this end, we generated a lung carcinoma cell line (A549) with constitutive ACE2 overexpression (A549^ACE2^). Once again, virapinib presented a dose-dependent effect in inhibiting the infection by SARS-CoV-2 and no detectable toxicity in A549^ACE2^ cells **(Fig. 4B, Fig. S2D)**. Furthermore, the compound reduced the incidence of multinucleated syncytia in A549^ACE2^ cells exposed to SARS-CoV-2 (**Fig. 4C, D**), which has been previously documented as a result of SARS-CoV-2 infection due to the interaction of membrane-present S protein with ACE2 of neighboring cells^9–11^. Importantly, and while we focused on virapinib for the general characterization reported in this study, structure-activity relationship (SAR) analyses revealed that there is significant room to improve the antiviral potency of this chemical series (**Fig. S3**)

After confirming the efficacy of virapinib against the ancestral strain, we tested the effects of the compound against different SARS-CoV-2 variants. Since its emergence, SARS-Cov-2 has been constantly mutating, and the World Health Organization (WHO) has classified several of these mutants as Variants of Concern (VOC), the most notable being Alpha (B.1.1.7), Delta (B.1.617.2), and Omicron sub-lineages (e.g. BA.1). Importantly, virapinib showed anti-infective activity against all variants in both Vero E6 and A549^ACE2^ cells, and the concentration of the compound needed to limit infection was correlated with the infectivity of each VOC (higher concentrations for the most infective strains) (**Fig. S4**).

Finally, we tested the efficacy of virapinib in a well-established liver spheroid model of extrapulmonary SARS-CoV-2 infection^12^. The model is extensively characterized and previously contributed to the identification of the antiviral activity of the JAK/STAT inhibitor baricitinib ^13,14^, which facilitated its rapid clinical translation and remains only one of four compounds endorsed by the WHO for severe Covid-19 ^15^. Notably, virapinib efficiently reduced SARS-CoV-2 infectivity in this model **(Fig. 4E)**, to a similar extent as remdesivir, a nucleoside analog that inhibits the viral RNA-dependent RNA polymerase^16^.

### Virapinib limits viral entry by inhibiting macropinocytosis

Having confirmed that virapinib can inhibit SARS-CoV-2 infection in several *in vitro* models, we next investigated its mechanism of action. Given that our screening assay was based on PV^s^ expressing the S protein and host cells expressing ACE2, we tested whether the compound affected these factors or their interaction. The compound did not affect the cellular distribution of ACE2 or the proteolytic processing of S (**Fig. S5A, B**). Moreover, in contrast to nanobodies targeting the S protein^17^, the binding between recombinant S and ACE2 was not affected by virapinib, as measured by biolayer interferometry (**Fig. S5C**). On the other hand, SARS-CoV-2 replication in A549^ACE2^ cells, which was efficiently inhibited by remdesivir, was not affected by virapinib, suggesting that the mechanism of action of the compounds is related to viral entry (**Fig. S5D**).

Next, we performed RNA sequencing (RNA-seq) in MCF-7 breast cancer cells treated with 10 μM virapinib for 6 h and analyzed differentially expressed genes (DEGs). Of note, the choice of the cell line, dose and times of treatment was done to be able to compare these data against the drug-associated transcriptional signatures existing at the Connectivity Map (CMap) database, which can provide hypothesis on the mechanism of action of a given drug on the basis of similarities between its transcriptional signature to that of other drugs^18^. Strikingly, most genes whose expression was significantly upregulated by virapinib were related to sterol biosynthesis, such as HMG-CoA synthetase and reductase (HMGCS and HMGCR), and the mevalonate kinase (MVK) (**Fig. 5A**). Accordingly, pathway enrichment analyses revealed “steroid biosynthesis” as the most significantly induced by virapinib, suggesting that the compound had an impact on membrane biology (**Fig. 5B**). RNA-seq data were validated by subsequent experiments, where we confirmed that virapinib treatment increased the levels of *HMGCR*, *HMGCS,* and *MVK* mRNAs (**Fig.5C**). As a control, we used methyl β-cyclodextrin (MβCD), which stimulates sterol biosynthesis by depleting cholesterol. Interestingly, however, virapinib did not affect cholesterol levels or its distribution (**Fig. 5D,E**).

**Figure 5.**
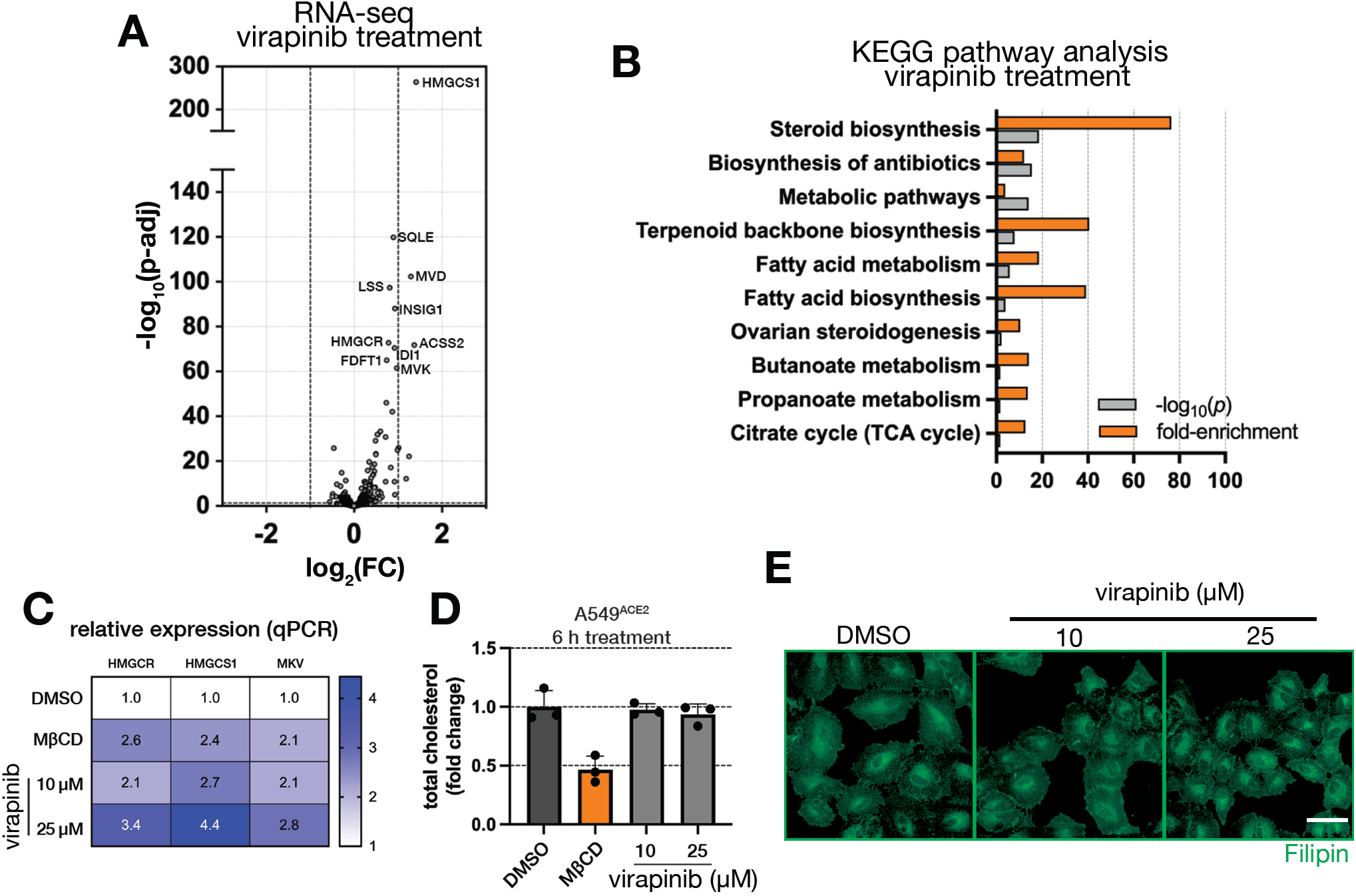
Virapinib affects steroid metabolism. (**A**) Volcano plot showing gene expression changes in MCF-7 cells treated with virapinib at 10 μM for 6 h. The cell line and dose were selected to enable the use of the obtained RNA-seq in databases of drug-associated transcriptional signatures to infer a potential mechanism of action. (**B**) Enrichment analysis of KEGG pathways upregulated in MCF-7 cells treated for 6 h with 10 μM virapinib. Genes with *p* < 0.05 and logFC > 1.2 were used as an input in the DAVID analysis tool. (**C**) RT-qPCR analysis of genes encoding enzymes from early steps of the cholesterol biosynthesis pathway. A549^ACE2^ cells were treated for 6 h with virapinib or methyl-ß-cyclodextrin (MβCD). Presented values are the means of two independent experiments, each with triplicates. (**D**) Representative data of quantification of cholesterol levels in A549^ACE2^ cells treated for 6 h with DMSO, 5 mM MβCD or virapinib. Analysis was done using a cholesterol Glo kit. Raw luminescence was normalized to the number of nuclei (assessed by live-cell microscopy prior cholesterol content analysis). Viability-corrected values are normalized to DMSO treated samples. (**E**) Representative microscopy images of A549 cells treated for 16 h with virapinib at the indicated doses and stained with Filipin (green) to visualize cholesterol levels. Scale bar (white) represents 50 μm.

In parallel to our RNA-seq experiments, we conducted immunofluorescence analyses of SARS-CoV-2 N protein at different timepoints post infection to evaluate how virapinib prevented viral infection. In control (DMSO) A549^ACE2^ cells, the staining pattern showed a clear development from small puncta at the periphery of the cell (1-2 h post infection (h.p.i.)), through a clear dissemination within the cell and the signal intensity increase at 4 h.p.i., to a pan-cytoplasmic staining at 6 h.p.i (**Fig. 6A**). In contrast, cells treated with virapinib showed a clear lag in the development of this pattern, indicating that either entry or trafficking of the virus was impaired (**Fig. 6A**). Quantification of this phenomenon by machine learning-based object classification confirmed our qualitative microscopic observations (**Fig. 6B**).

**Figure 6.**
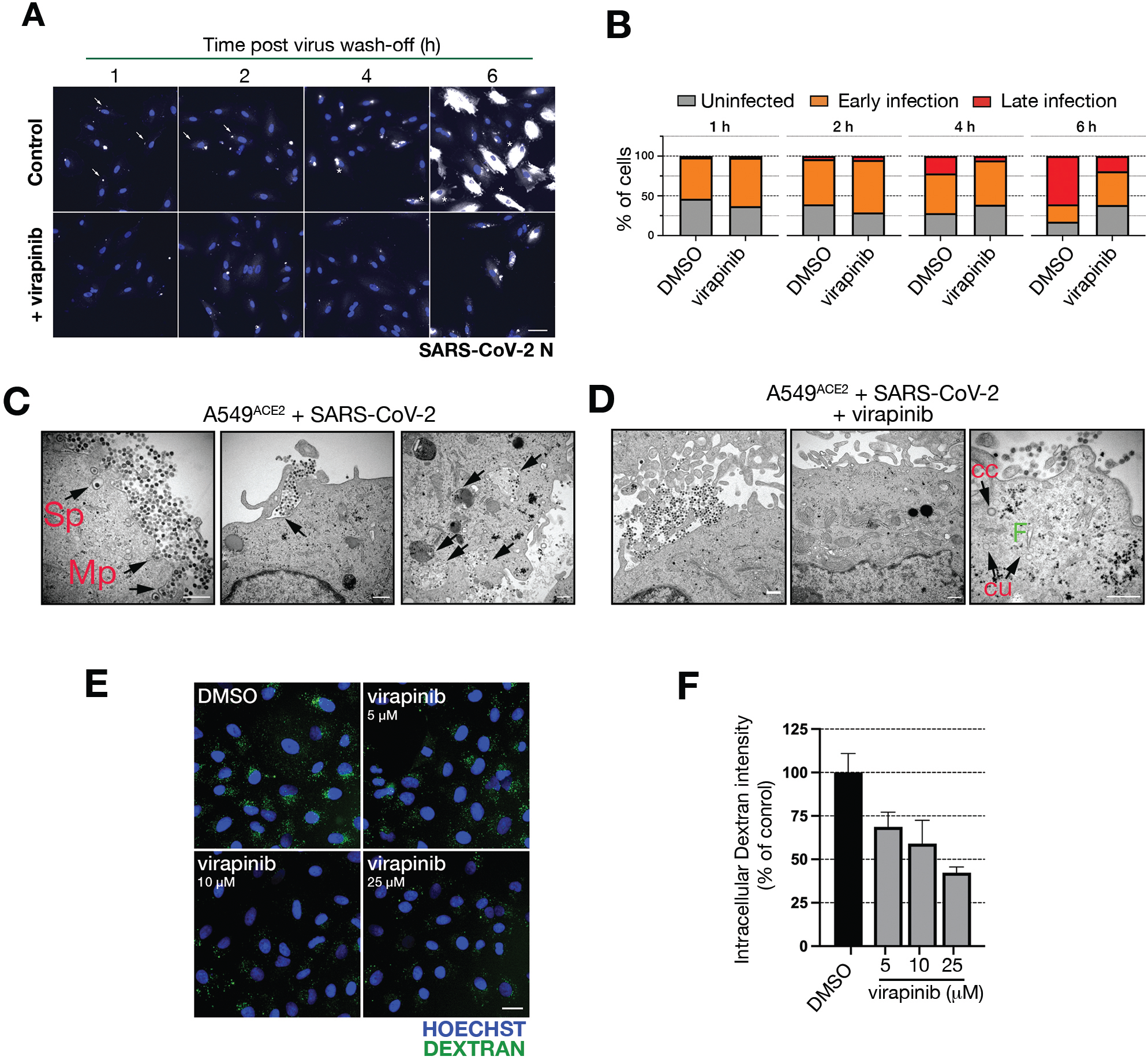
Virapinib inhibits viral entry by macropinocytosis. (**A**) Time-resolved monitoring of SARS-Cov-2 entry and dissemination. A549^ACE2^ cells were treated with DMSO or 10 μM virapinib for 16 h prior to 1 h infection. Cells were fixed at 1, 2, 4, and 6 h after virus wash-off, and stained for the detection of SARS-CoV-2 N protein (white) as in previous experiments. Arrows indicates cells classified as ‘early infection’, while asterisks indicate ‘late infection’. (**B**) Quantification of (C). Graphs show the percentage of each phenotype at a given time-point. Object classification was done by machine learning using CellProfiler Analyst and 300-500 cells were analysed for each sample. (**C**) Representative TEM images of A549^ACE2^ cells infected with SARS-Cov-2. [Left] Image illustrating the 2 modes of viral entry (single particle entry (Sp) and macropinocytosis (Mp). [Middle] Image illustrating the initial steps of macropinocytosis. [Right] Examples of the different steps of the macropinocytosis cycle, from early invaginations to internal autophagosomes at various stages of maturation. Scale bar (white) indicates 500 nm. (**D**) Representative TEM images of A549^ACE2^ cells infected with SARS-Cov-2 and treated with 25 μM virapinib. Virapinib was added for 2 h before fixation, following infection with SARS-CoV-2 for 1 h. [Left] The accumulation of viral particles at the cell surface did not induce their internalization in virapinib treated cells, although membrane ruffled or protrusions normally formed in them. [Middle] The ultrastructural organization of the membrane and of various cytoplasmic organelles such as mitochondria, endoplasmic reticulum, Golgi complex, endosomes and endocytic vesicles of micropinocytosis is unaffected by the drug. [Right] Virapinib treated cells show normal looking clathrin-coated (cc) or uncoated (cu) endocytic vesicles from micropinocytosis and their fusion (F) to early endosomes. Scale bar (white) indicates 500 nm. (**D**) Representative images from a macropinocytosis assay based on the uptake of a fluorescent high-molecular weight (HMW) dextran (green). A549 cells were treated for 16 h prior to a 2 h incubation with the dextran. The graph on the right shows the quantification of the assay. Each data point represents the mean of 1 well in which 1500-2000 cells were analysed.

Given that our transcriptomic analyses indicated that the effect of virapinib could be related to viral entry and lipid metabolism, we next analyzed the effect of the compound by transmission electron microscopy (TEM), particularly focusing on membrane dynamics, in A549^ACE2^ cells exposed to SARS-CoV-2. In this experimental setup, viral particles infected host cells through two distinct entry routes: endocytosis of individual viruses or bulk entry by macropinocytosis, a known mechanism of viral infection^19^, whereby numerous virus particles are engulfed together with extracellular material into irregularly shaped large vesicles (**Fig. 6C**). Importantly, macropinocytosis was by large the predominant mechanism for SARS-CoV-2 entry. Furthermore, qualitative analysis of TEM images indicated that virapinib selectively inhibited the macropinocytic route, while not having a detectable impact on single-particle entry, the cell membrane or the integrity of intracellular organelles (**Fig. 6D**). To directly test the effect of virapinib on macropinocytosis, we took advantage of an established assay based on the uptake of a fluorescent high-molecular-weight dextran^20^, which revealed that virapinib limited dextran uptake in A549 cells (**Fig. 6D**). Together, these experiments show that virapinib inhibits macropinocytosis, which is a major mechanism of SARS-Cov-2 entry into host cells.

### Virapinib inhibits infection by Mpox, TBEV and Ebola viruses

Given that macropinocytosis has been reported to be the preferred mode of entry for many viruses, we finally explored whether virapinib could have broader antiviral activity. To do this, we tested its effect on the infection with the following viruses: Andes virus (ANDV; *Hantaviridae* family; enveloped; ssRNA), Dengue virus (DENV; *Flaviviridae* family; enveloped; ssRNA), Mpox (MPXV; *Poxviridae* family; enveloped; dsDNA), Tick-borne encephalitis virus (TBEV; *Flaviviridae* family; enveloped; ssRNA), a EGFP-expressing adenovirus (*Adenoviridae* family, non-enveloped, dsDNA) and a VSV-pseudotyped lentivirus expressing the envelope glycoprotein from Ebola virus. Virapinib showed dose-dependent antiviral activity against TBEV, Monkeypox virus, and Ebola-pseudotyped VSV (**Fig. 7A**). We did not observe any decrease in the infection rate of ANDV (**Fig. 7B**) and only a small effect on adenoviral infection (**Fig. 7C**). Interestingly, there was a dose-dependent increase in the infection rate with DENV (**Fig. 7D**), potentially reflecting that the inhibition of macropinocytosis could enhance other endocytic routes.

**Figure 7.**
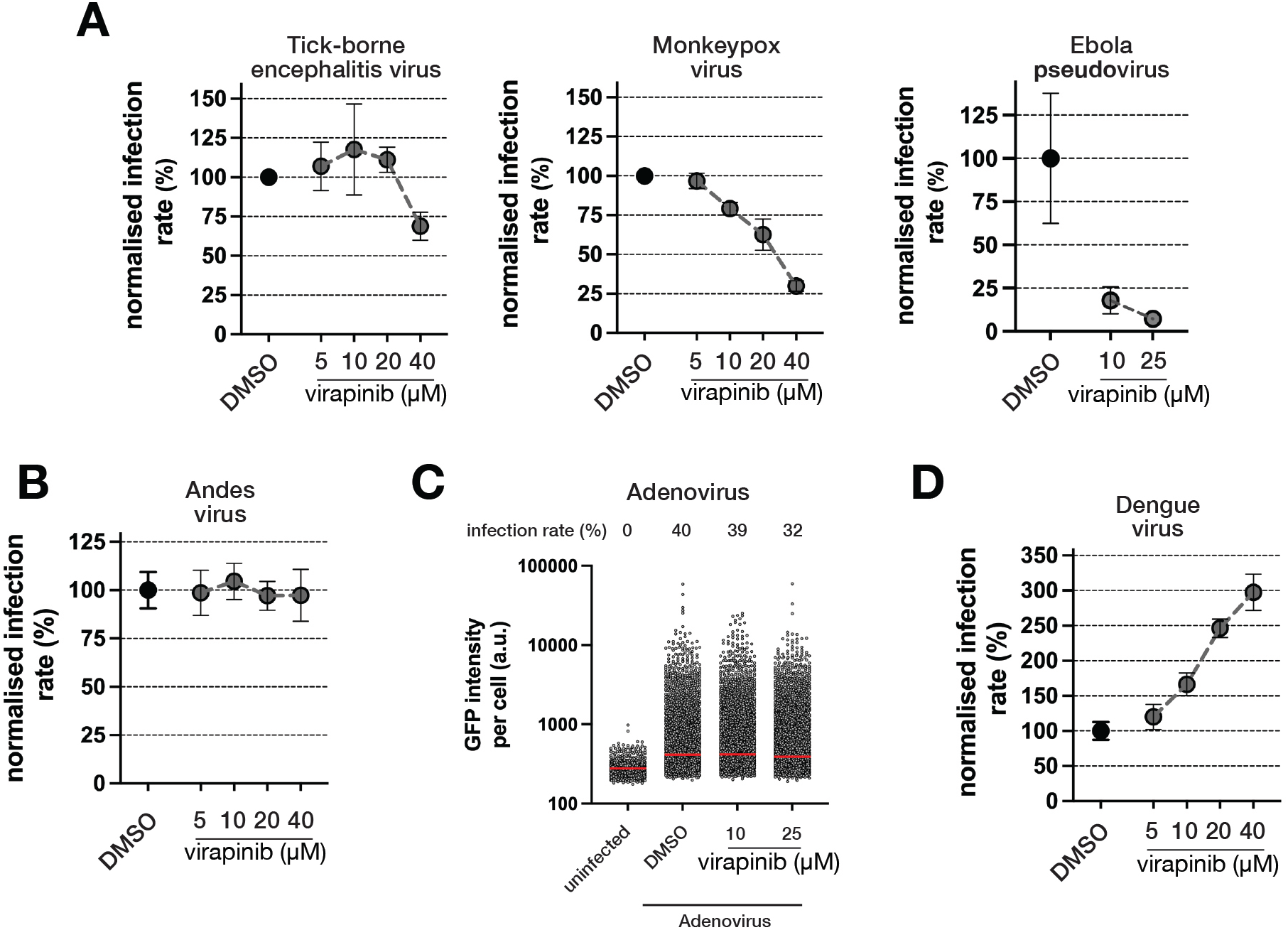
Virapinib is a broad-spectrum antiviral. (**A**) Virapinib has antiviral activity against viruses from different families. A549 cells were pre-treated for 6 h prior to the addition of the indicated virus. Each data point represents the mean ± S.D. of 3 wells, each containing ∼15000 cells. (**B**) As in (A), but with Andes virus, where virapinib showed no effect. (**C**) Activity of virapinib against infection of MEF with an adenovirus expressing a Cre-EGFP fusion protein. MEF expressing CAR were treated for 16 h prior to a 4 h infection. Cells were imaged 24 h post-infection. The panel represents the HTM-dependent quantification of GFP expression. Each data point represents one cell. Red lines mark median values. (**D**) As in (A,B), but with Dengue virus where surprisingly virapinib increases infection rates.

## DISCUSSION

The onset of the Covid-19 pandemic triggered a world-wide initiative to find strategies that could limit the spread and severity of the disease^21^. To a large extent, this was achieved by a combination of collective vaccination and the use of anti-inflammatory treatments to treat the fatal cytokine storm triggered by the disease. In what regards to chemical inhibitors of viral infection, three drugs have received approval for their medical use (paxlovid, remdesivir and molnupiravir). Of note, these approaches target viral proteins, and no host-targeting antiviral has been approved for the treatment of Covid-19. While intense research efforts were placed on drug-repurposing campaigns, identified hits did not hold in the clinical setting^22^.

The difficulty of antiviral development is not limited to SARS-CoV-2 and, currently, only a fraction of pathogenic human viruses can be treated with antiviral drugs. This is partly due to the high demands for safety to work with human pathogens^23^, which to some extent is solved by the use of non-replicating pseudotype viruses. This approach models the initial infection step of live viruses, and was used as a platform on a drug-repurposing screen for inhibitors of SARS-CoV-2 entry^24^. Given the strong focus on repurposing by many other laboratories, we decided to explore another chemical space hoping to find new chemical entities showing anti-infection properties. The discovery of virapinib further validates the usefulness of this approach as a discovery platform for antiviral candidates that limit the entry step. Our subsequent experiments revealed that the compound specifically limited SARS-CoV-2 entry by macropinocytosis. Of note, published studies had reported conflicting views about SARS-CoV-2 mode of entry, with some reporting clathrin-dependent endocytosis as the preferred route^25^ while others implicating macropinocytosis^26^. Our TEM analysis was clarifying in this regard, illustrating how SARS-CoV-2 can simultaneously use both entry pathways, yet only macropinocytosis is inhibited by virapinib.

As to how virapinib regulates macropinocytosis, this remains to be fully understood. It is remarkable that our transcriptomic analyses revealed that the compound triggered very subtle changes on gene expression, consistent with the fact that the compound did not have any obvious phenotypic impact on host cells. Furthermore, the observation that the few genes which expression was altered by virapinib were mostly related to steroid biosynthesis suggests that the compound might affect interactions happening at the lipid bilayer membrane. Along these lines, many studies have implicated cholesterol in SARS-CoV-2 infection^27–31^. Accordingly, decreasing cholesterol with methyl-ß-cyclodextrin inhibits infection with SARS-CoV-2, while supplementation with exogenous cholesterol potentiates the infection^29,32^. Since virapinib does not affect cholesterol levels or distribution, we suggest that it might integrate at membranes and disrupt their physical properties limiting the formation of processes like membrane ruffling, which are necessary for macropinocytosis. Our TEM images provide indications that this is the case. Further supporting this hypothesis, the chemical structure of virapinib contains a fatty-acid-like moiety, which should facilitate membrane-targeting of the compound. Further studies of membrane biology and biophysics should be clarifying to better understand how virapinib impairs macropinocytosis.

Even if our original aim was to find antiviral candidates for SARS-CoV-2, the fact that the compound inhibits macropinocytosis extends its potential usefulness for other pathologies. Given that macropinocytosis is used by multiple viruses as a mechanism for cell entry, it would be worthwhile to systematically test its antiviral capacity against a wide range of viruses. Our initial analysis with a limited set of viruses supports that the compound might be efficacious against multiple pathogens. In addition, macropinocytosis has been shown to have tumor promoting roles (reviewed in ^33^), as well as to contribute to the cell-to-cell propagation of toxic protein aggregates in neurodegenerative diseases^34^. Thus, and besides its potential usefulness as an antiviral, our work here provides a new tool for the scientific community that should help to address the contribution of macropinocytosis to disease, and potentially help to mitigate the severity of these pathologies.

To end, we must clearly state that the work reported here is to be understood as an exercise of basic research, which highlights the usefulness of focusing the chemical screens for antivirals on libraries other than drug-repurposing ones, and supports the concept of targeting endocytic pathways for limiting viral infections. Even if initial SAR indicates that the potency of virapinib can be substantially enhanced, additional work on this chemical series, including in vivo experiments, would be needed to better define its antiviral efficacy.

## MATERIALS AND METHODS

### Cell lines

HEK293T (ATCC CRL-11268) and A549 (ATCC CCL-185) cells were grown in Dulbecco’s Modified Eagle Medium (Gibco™, 31966021) supplemented with 10% Fetal Bovine Serum (Sigma-Aldrich, F7524) and 1% penicillin-streptomycin (Gibco™, 15140122). Vero E6 (ATCC CRL-1586) cells were grown in Minimum Essential medium supplemented with 7.5% FBS, HEPES, L-Glutamine, 100 U/ml penicillin and 100 mg/ml streptomycin. All cell lines were grown at 37 °C, 5% oxygen and 5% carbon dioxide; cells were routinely passaged before reaching confluency. HEK 293T^ACE2^ and A549^ACE2^ cells were generated by a transduction with a lentivirus expressing human ACE2 (kind gift from Benjamin Murrel, Karolinska Institutet) in the presence of 6 μg/ml Polybrene (Sigma-Aldrich, H9268). Clones were isolated and selected for homogeneous ACE2 expression as determined by HTM. HEK 293T cells expressing human-codon optimized S Delta/B.1.617.2 (cDNA kindly provided by the G2P-UK National Virology Consortium and the Barclay Lab at Imperial College London) in a doxycycline(dox)-dependent manner were generated by CRISPR-Cas9-directed insertion of a Tet-On® 3G tet-repressor into the human Rosa locus, and the S cDNA under the TRE3GS promoter. Plasmids carrying the transgene and Cas9/sgRNA were co-transfected using Lipofectamine 3000 according to the manufacturer’s protocol. Clones were selected using Zeocin™ (Gibco™, R25001).

### Pseudotype viral production

S- and VSVg-enveloped lentiviruses were produced in HEK 293T cells. To do so, cells were co-transfected with enveloping (pCMV14-3X-Flag-SARS-CoV-2 S (Addgene #145780) for PV^s^ or pMD2.G (Addgene #12259) for VSVg-PV), packaging (psPAX2, Addgene #12260), and transfer (pHR-CMV-GFP, Addgene #14858) plasmids. Plasmids were mixed at 2:1:1 molar ratio and transfected using Lipofectamine 2000 according to manufacturer’s protocol. Media was exchanged after 16h, and virus-rich supernatants harvested after 24 and 48 additional hours. An aliquot of the PV^s^ preparation was boiled with Laemmli buffer and analysed by WB to detect the presence of the S protein. Pseudotype viruses for Ebola were produced in the same manner, using an enveloping plasmid expressing the Ebola glycoprotein cDNA (kind gift from Jochen Bodem). Virus-rich supernatants were concentrated using Lenti- X™ Concentrator (Takara Bio, 631231) according to the manufacturer’s protocol.

### Western blotting

Cells were washed with PBS, scraped on ice into RIPA buffer (Thermo Scientific, PI-89901), sonicated, and centrifuged at 14’000 x g for 15 min at 4 °C. Supernatant was recover and protein concentration was measured using the DC™ Protein Assay Kit II (Bio-Rad, 5000112). Lysates were boiled in the NuPAGE™ LDS Sample Buffer (Invitrogen™, NP0007) supplemented with NuPAGE™ Reducing Agent (Invitrogen™, NP0009). Samples were electrophoresed on a Bis-Tris gel, and electroblotted onto a nitrocellulose membrane (Bio-Rad, 1704270). Membranes were blocked in 5% milk/TBS-Tween buffer and incubated overnight at 4 °C with primary antibodies. On the following day, membranes were washed in TBS-Tween, incubated with secondary antibody coupled to horseradish peroxidase for 1h at RT, and washed again in TBS-Tween. Signal was developed using a kit (Thermo Scientific, 10220294) and detected in an Amersham™ Imager 600 (GE Healthcare). A full list of the antibodies used in this study is available in **Table S1**. The antibody used for the detection of SARS-CoV-2 N protein was a kind gift from Pamela Österlund^35^.

### Microscopy

To prepare samples for fluorescence microscopy, cells were fixed with 4% formaldehyde for 15 min at RT, followed by a 10-min permeabilization with 0.1% Triton X-100, a 30-min blocking with blocking buffer (3% BSA/0.1% Tween-20/PBS), overnight incubation with primary antibodies at 4 °C, and a 1h incubation with secondary antibody solution supplemented with 2 μM Hoechst 3342 (Thermo Scientific™, 62249) and 1 μM CellTracker™ Orange CMTMR Dye (Thermo Scientific™, C2927) or Alexa Fluor™ Plus 555 Phalloidin (Thermo Scientific™, A30106). Image acquisition was done using an IN Cell Analyzer 2200 (GE Healthcare) high-throughput microscope. Quantitative image analysis was performed using CellProfiler^36^, and the automated classification of viral entry phenotypes by machine learning was done using CellProfiler Analyst^37^.

### Chemical screen

For the primary screen, 293T^ACE2^ cells were seeded onto 384 well plates (BD Falcon, 353962) and incubated overnight. On the following day, chemicals from a library of 1008 compounds designed for the discovery of preventing protein-protein interactions (PPi library, Asinex, kindly provided by the Chemical Biology Consortium Sweden) were added to cells 6h prior to the addition of a PV^s^ solution containing polybrene. The final concentration of the compounds was 25 μM and E-64D (Tocris, 4545) was used in some of the wells as a control for infection inhibition. After 72h, cells were fixed for 15 min with 4% formaldehyde in the presence of 2 μM Hoechst 3342. After a was in PBS, plates were sealed, and samples imaged by HTM. Image analysis was done in CellProfiler^36^ and data processing was done using the KNIME Analytics Platform (Knime) and Excel (Microsoft) software. All statistical analyses in the manuscript were performed using Prism software (GraphPad Software) using the indicated tests for each experiment.

### Viral infections

Cells were seeded on the day before infection. Infection was done by replacing growth media with virus solutions, followed by a 1h incubation, PBS wash, and re-addition of full growth medium. Cells were fixed for 24 h (48 h for Dengue virus) after infection with 4% formaldehyde for 15 min at RT. Afterwards, cells were washed once with PBS and the plate was placed in 70% ethanol prior to its removal from the BSL-3 lab. Detection of infected cells was done by measuring the expression of viral proteins by HTM as defined above. All virus-rich supernatants were ultracentrifuged at 45’000 x g for 4 h and titrated via an end-point dilution assay in Vero E6 cells. The TCID50 (50% tissue culture infection dose) per milliliter was calculated using the Spearman-Karber method. Infection with the Ebola pseudovirus and calculation of infection rates was done as for PV^s^, but in A549 cells. Infection with Cre-EGFP expressing adenovirus (Vector Biolabs, 1700) was done in mouse embryonic fibroblasts expressing the CAR receptor that were pre-treated with compounds for 16 h. The infection was done at a MOI of 20 for 4h. 24h after infection, cells were fixed and imaged with an Opera Phenix HTM (Perkin Elmer), and infection evaluated on the basis of EGFP expression.

### Liver spheroids

3D primary human hepatocyte (PHH) spheroids were generated and cultured as previously described^38^. In short, PHH were seeded at a density of 1,500 cells per well in 96-well ultra-low attachment (ULA) plates and spheroids formed over the course of six days in PHH culture medium (Williams E medium supplemented with 2 mM L-glutamine, 100 units/ml penicillin, 100 μg/ml streptomycin, 10 μg/ml insulin, 5.5 μg/ml transferrin, 6.7 ng/ml sodium selenite, 100 nM dexamethasone) supplemented with 10% fetal bovine serum. After spheroids were formed, the medium was changed to serum-free PHH culture medium.

For viral infection, PHH spheroids were exposed overnight to the indicated concentrations of remdesivir or virapinib in serum-free PHH culture medium. Spheroids were then infected with SARS-CoV-2 (hCoV-19/sweden720-53846/2020) at a multiplicity of infection of 1 for 24 h. After 24 h, four spheroids per condition were pooled and washed 3 times with PBS before lysis. RNA was extracted and purified using the Quick-RNA Microprep Kit (Zymo Research Irvine, CA, USA) and relative levels of viral RNA were determined by qRT-PCR as previously described^39^. hCoV-19/sweden720-53846/2020 viruses were kindly provided by The Public Health Agency of Sweden to improve the quality of diagnostics relevant for infectious disease control, treatment and/or other studies of relevance for public health. A full list of viruses used in this study is available in **Table S1**.

### Transmission electron microscopy

A549^ACE2^ cells infected with SARS-CoV-2 (MOI = 10) for 1 h, and treated or not with virapinib were washed twice with PBS, and supplemented with fresh culture medium. After 2 h, cells were fixed in 2.5% glutaraldehyde in 0.1 M phosphate buffer pH 7.4 at RT for 1h followed by storage at +4°C. After fixation, cells were rinsed in 0.1M phosphate buffer and postfixed in 2% osmium tetroxide in 0.1 M phosphate buffer, pH 7.4 at 4 °C for 2 h. Following stepwise dehydration in ethanol and acetone, cells were embedded in LX-112 resin (Ladd Research). Ultrathin sections (∼80-100 nm) were prepared using an EM UC7 (Leica), contrasted with uranyl acetate followed by lead citrate, and examined in a HT7700 transmission electron microscope (Hitachi High-Technologies,) at 80 kV. Digital images were acquired using a 2kx2k Veleta CCD camera (Olympus Soft Imaging Solutions).

### RNA-seq

Total RNA was extracted from cell pellets using a Purelink RNA Mini Kit (Invitrogen #12183025) following manufacturer’s instructions. Total RNA was subjected to quality control with an Agilent Tapestation (#G2991BA). To construct libraries suitable for Illumina sequencing, an Illumina stranded mRNA prep ligation sample preparation protocol was used with a starting concentration of total RNA between 25-1000 ng. The protocol includes mRNA isolation, cDNA synthesis, ligation of adapters and amplification of indexed libraries. The yield and quality of the amplified libraries was analyzed using a Qubit by Thermo Fisher and the quality of the library was checked using the Agilent Tapestation. Indexed cDNA libraries were normalized and combined, and pools were sequenced using an Illumina platform. STAR^40^ was used for sequence alignment based on the GRCh38 DNA primary assembly reference build, and quantification was done using featureCounts^41^. Differential expression analyses were performed using DESeq2^42^. Generalized linear model (GLM) was fitted to the expression data and shrunken log2fold-change (LFC) using adaptive Student’s t prior shrinkage estimator^42,43^. Multiple testing correction was done using Benjamini-Hochberg (BH) method. Enrichment analyses on “Kyoto Encyclopedia of Genes and Genomes” (KEGG) Pathways enrichment analysis was done using DAVID^44^.

### Quantitative RT-PCR

Total RNA was extracted from cell pellets using a Purelink RNA Mini Kit (Invitrogen #12183025) following manufacturer’s instructions. Reverse transcription and PCR amplification were performed using TaqMan RNA-to-CT 1-Step Kit and the StepOnePlus™ Real-Time PCR Instrument (Applied Biosystems, Fisher Scientific). All primers used in this study are listed in **Table S1**.

### Cholesterol measurements

To visualize the cellular distribution of cholesterol, cells were fixed with methanol-free formaldehyde, washed with PBS, and stained with 10 µg/ml Filipin (Sigma-Aldrich, SAE0087) for 1 h in dark. After a PBS wash, samples were imaged with an InCell Analyzer 2200 HTM (GE Healthcare). Quantification of cellular cholesterol was done using the Cholesterol/Cholesterol Ester-Glo™ kit (Promega, J3190), according to the manufacturer’s protocol miniaturized to a 384-well plate format. Prior to this procedure, cells were stained live with Hoechst 3342 and nuclei were imaged. Cell numbers were then quantified using CellProfiler^36^ and used to normalize the luminescence signal from cholesterol measurements.

### Bio-layer interferometry

The kinetics of S protein binding to hACE2 was analyzed using bio-layer interferometry, using recombinant His-tagged and biotinylated human ACE2 protein (Sino Biological, Cat 10108-H08H-B) and recombinant SARS-Cov-2 His-tagged S protein (Sino Biological, Cat #40591-V08H). Binding was analyzed using an Octet RED96 instrument (ForteBio). Streptavidin-coated sensors were equilibrated in 10x kinetics buffer (1% w/v BSA, 0.2% Tween-20 in PBS) for 15 min at room temperature. Binding assays were conducted at 30℃. First, a baseline was established in 10x kinetics buffer for 120s. Biotinylated human ACE2 (2.5 ug/mL in 10x kinetics buffer) was immobilized on the sensors for 120s. The sensors were washed in 10x kinetics for 120s. Sensors were then dipped into recombinant SARS-Cov-2 S protein (a titration ranging from 199 nM to 6.2 nM) either alone, or in the presence of DMSO, or 1 μM nanobody, or 50 μM Virolin. All proteins were diluted in 10x kinetics buffer. The analytes were allowed to associate for 180s, followed by dissociation in 10x kinetics for 180s. Data was analyzed using Octet ForteBio Analysis 9.0 software. KD values were measured by subtracting a reference sensor (ACE2 loaded, but no analyte) from all curves, then fit using a 1:1 model. An anti-Spike nanobody (kind gift from Benjamin Murrel) was used as a control for an interaction inhibitor^17^.

### SARS-CoV-2 S protein processing

HEK 293T cells expressing SARS-CoV-2 S Delta protein in a dox-inducible manner were seeded 24 h before treatment with DMSO, Virapinib (25 μM), the Furin inhibitor Decanoyl-RVKR-CMK (50 μM; Tocris, 3501) or the TMPRSS2 inhibitor Camostat Mesylate (50 μM, Sigma-Aldrich, SML0057). After 6 h, expression of the S protein was induced by the addition of 50 ng/ml dox (Sigma-Aldrich, D9891) for 24 h. Cells were harvested after additional 24 h and analysed by WB.

## Data Availability

RNA-seq data associated to this study are available at the GEO repository with accession number GSE246407.

## ACKNOWLEDGEMENTS

We thank the Chemical Biology Consortium Sweden for their help with the chemical libraries; Dr Lars Haag at the Electron Microscopy Unit, Karolinska Institutet for support with TEM experiments and the Bioinformatics and Expression Analysis facility, Karolinska Institutet, for help with RNA-seq. Research was funded by grants from the Cancerfonden foundation (CAN 21/1529) and the Swedish Research Council (VR) (538-2014-31) to OF; the Swedish Society for Medical Research (P18-0098; PD20-0153) to S.C.W.; the Swedish Research Council (grant agreement numbers 2019-01837 and 2021-02801) and the EU/EFPIA/OICR/McGill/KTH/Diamond Innovative Medicines Initiative 2 Joint Undertaking (EUbOPEN grant number 875510) to V.M.L.; the Swedish Research Council Starting Grant (2020-02682) and Human Frontier Science Program (RGP0025/2022) to E.S.; and a Marie Skłodowska-Curie Actions Postdoctoral Fellowship (101059180) and a KI Virus Research Grant to J.S.

## AUTHOR CONTRIBUTIONS

B.P. contributed to most experiments and data analyses, preparation of the figures and to the writing of the manuscript. W.C. and J.K. contributed to experiments of viral infections. A.C. participated in the study design and initial screen. M.HA. provided help with the chemistry. M.B. helped in macropinocytosis experiments. L.L. provided technical help. M.H. contributed to the chemical screen. J.I. and G.B. performed S-ACE2 interaction assays. S. M.M. contributed to adenoviral infection experiments. J.S. and E.S. helped with S-ACE2 interaction analyses. S.W. and V.M.L. contributed the experiments in liver spheroids. M.L. helped with TEM analyses. D.H. helped to coordinate the study. O.F-C. supervised the study and wrote the MS.

