## Supplementary material for "Discovery of a novel inhibitor of macropinocytosis with antiviral activity": Figures S1-S5

Porebski *et al.* **Fig S1**

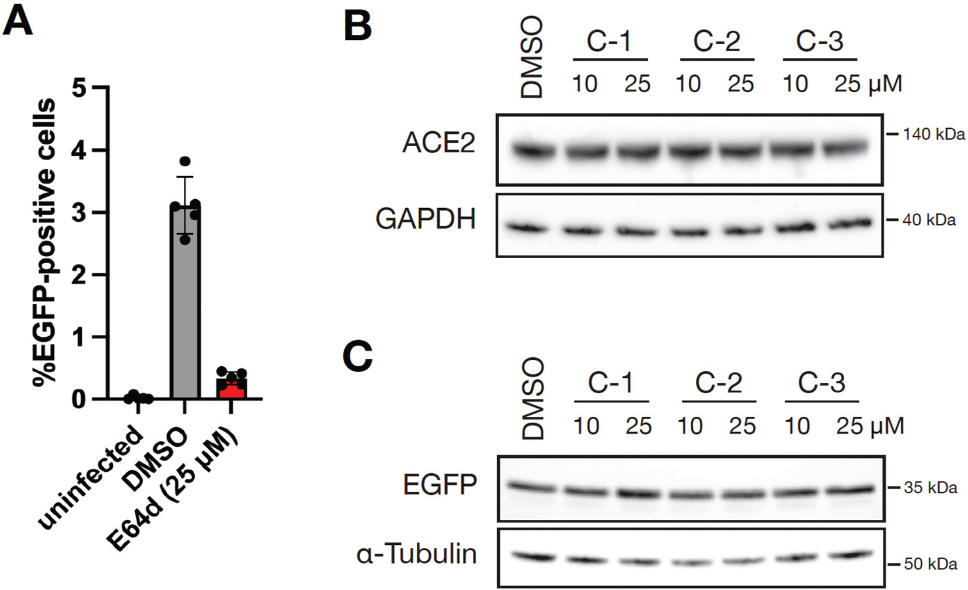

**Figure S1. Additional data on the chemical screen on PV$^{s}$ infection.** (**A**) Raw infection rate data from the screen as calculated on the basis of GFP expressing cells. Each data point represents the mean infection rate within one plate, calculated from 8 separate wells. Note the significant effect of E-64d in preventing PV$^{s}$ infection, providing a good assay window for the screen. (**B**) WB analysis of the effect of the indicated compounds (72 h) on ACE2 protein levels in HEK 293T$^{ACE2}$ cells. GAPDH levels are shown as a loading control. (**C**) Analysis of the effect of the indicated compounds (72 h) on EGFP protein levels in PV$^{s}$ infected HEK 293T$^{ACE2}$ cells. $\alpha$-TUBULIN levels are shown as a loading control.

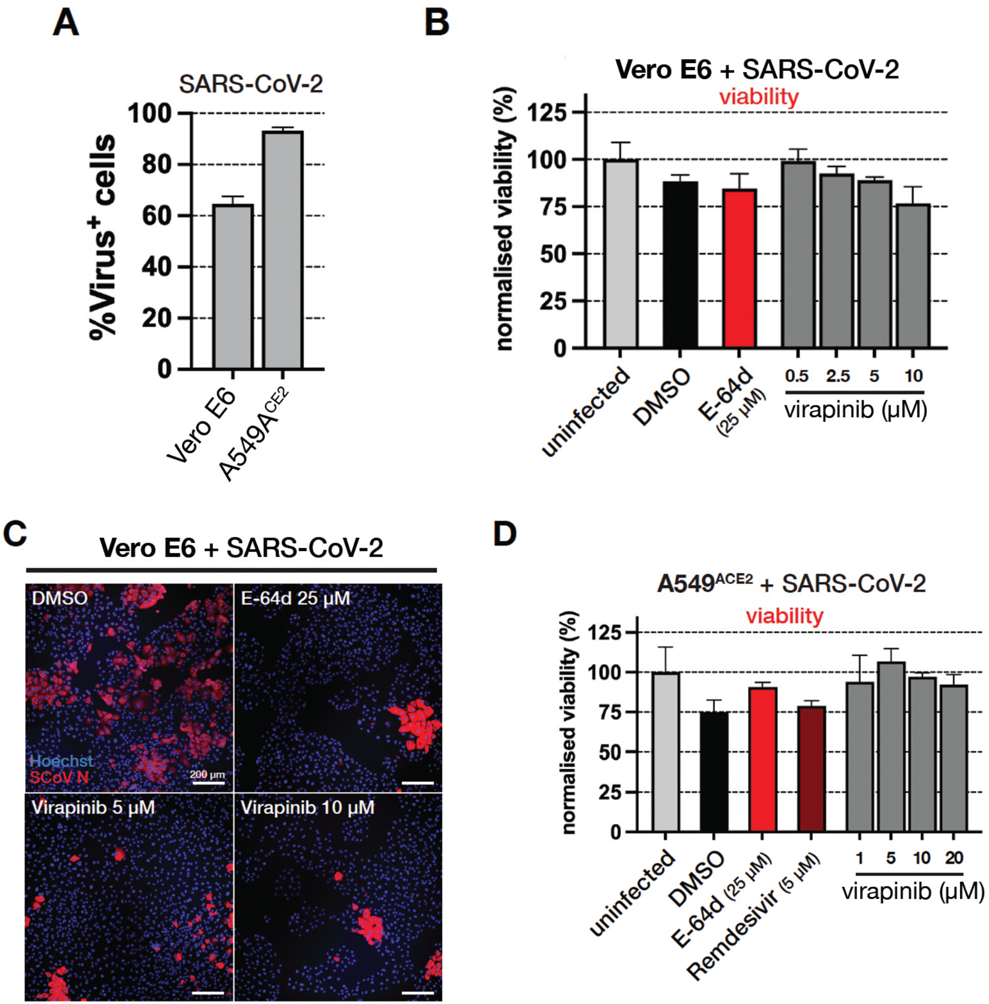
A
SARS-CoV-2
%Virus+ cells
100
80
60
40
20
0
Vero E6
A549ACE2
B
Vero E6 + SARS-CoV-2
viability
normalised viability (%)
125
100
75
50
25
0
uninfected
DMSO
E-64d (25 µM)
0.5
2.5
5
10
virapinib (µM)
C
Vero E6 + SARS-CoV-2
DMSO
E-64d 25 µM
Hoechst
SCoV N
200 µm
Virapinib 5 µM
Virapinib 10 µM
D
A549ACE2 + SARS-CoV-2
viability
normalised viability (%)
125
100
75
50
25
0
uninfected
DMSO
E-64d (25 µM)
Remdesivir (5 µM)
1
5
10
20
virapinib (µM)

**Figure S2. Effects of virapinib in SARS-CoV-2 infection and cell viability.** (**A**) Raw infection rate of SARS-CoV-2 in Vero E6 and A549$^{ACE2}$ cells. Plotted data represent the mean percentage of Virus-positive cells from 3 independent samples. In each replicate, 5000-10000 cells were imaged and quantified. Infected cells were detected on the basis of SARS-CoV-2 N protein expression. (**B**) Viability of Vero E6 cells treated with virapinib. Viability was assessed by counting nuclei stained with Hoechst 33342 by HTM. Plotted data represent the mean percentage of nuclei for each condition from 3 replicates. In each replicate, 5000-10000 cells were imaged and quantified. (**C**) Representative images of Vero E6 cells treated with vehicle (DMSO), E-64d or virapinib upon SARS-CoV-2 infection. Infection was detected on the basis of cells expressing the N protein from SARS-CoV-2 (red). Hoechst 33342 (blue) was used to visualize nuclei. Scale bar (white) represents 200 μm. (**D**) As in (B), but in A549$^{ACE2}$ cells.

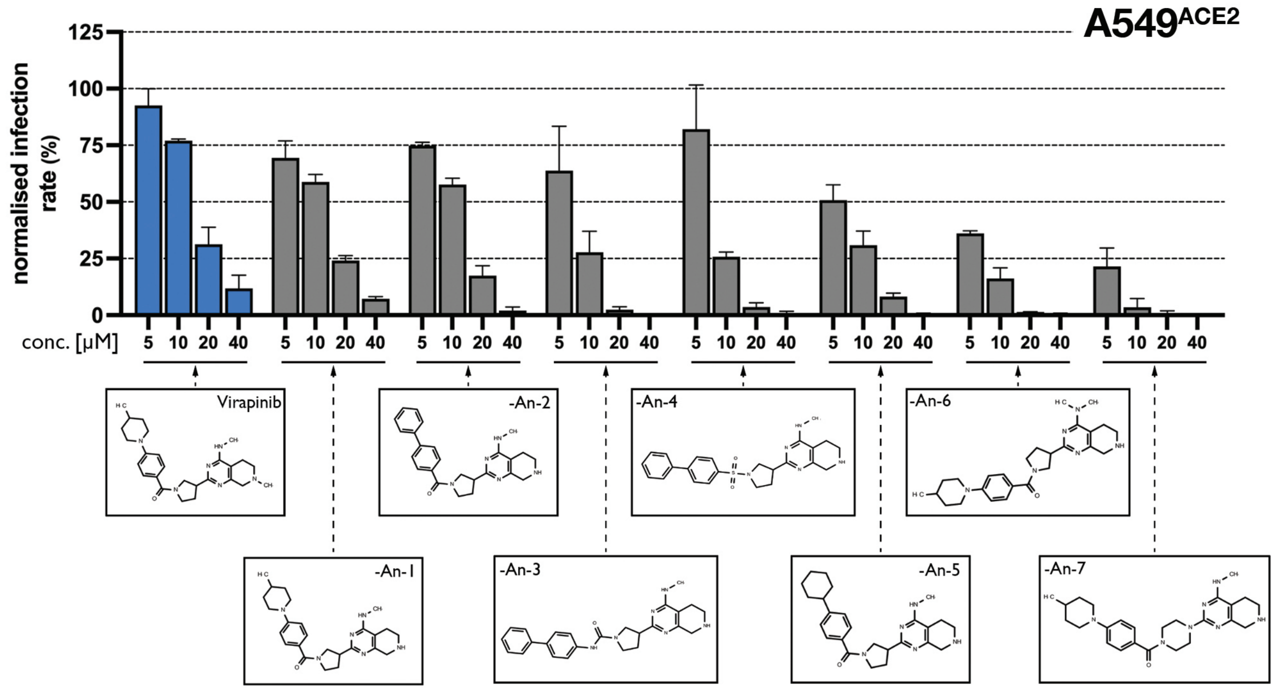
A549ACE2
125
100
75
50
25
0
normalised infection rate (%)
conc. [µM]
5 10 20 40
5 10 20 40
5 10 20 40
5 10 20 40
5 10 20 40
5 10 20 40
5 10 20 40
5 10 20 40
Virapinib
-An-1
-An-2
-An-3
-An-4
-An-5
-An-6
-An-7

**Figure S3. SAR identifies virapinib analogues with increased anti-infective activity.** Dose-response of virapinib and 7 analogues defined (An-1/7) by SAR studies against ancestral SARS-CoV-2 infection in A549$^{ACE2}$ cells. Cells were pre-treated with the compound for 6 h, followed by viral infection. Cells were fixed at 24hpi and infection rates were calculated by HTM quantifying cells expressing the N-protein from SARS-CoV-2. Each data point represents the mean ± S.D. of 3 wells, each containing ~15000 cells. The structure of each analogue is provided in a connected box.

Porebski *et al.* **Fig S4**

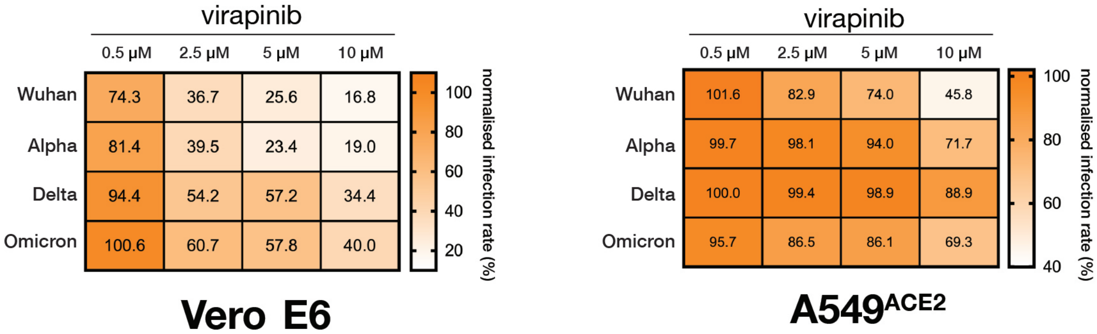

**Figure S4. Effect of virapinib against different SARS-CoV-2 variants.** Heatmaps illustrating infection rate of SARS-CoV-2 variants in Vero E6 cells (left) or $A549^{ACE2}$ (right) cells. Infection was performed and quantified as in **Fig. 4**. Each value represents the mean of 3 wells, each containing ~15000 cells.

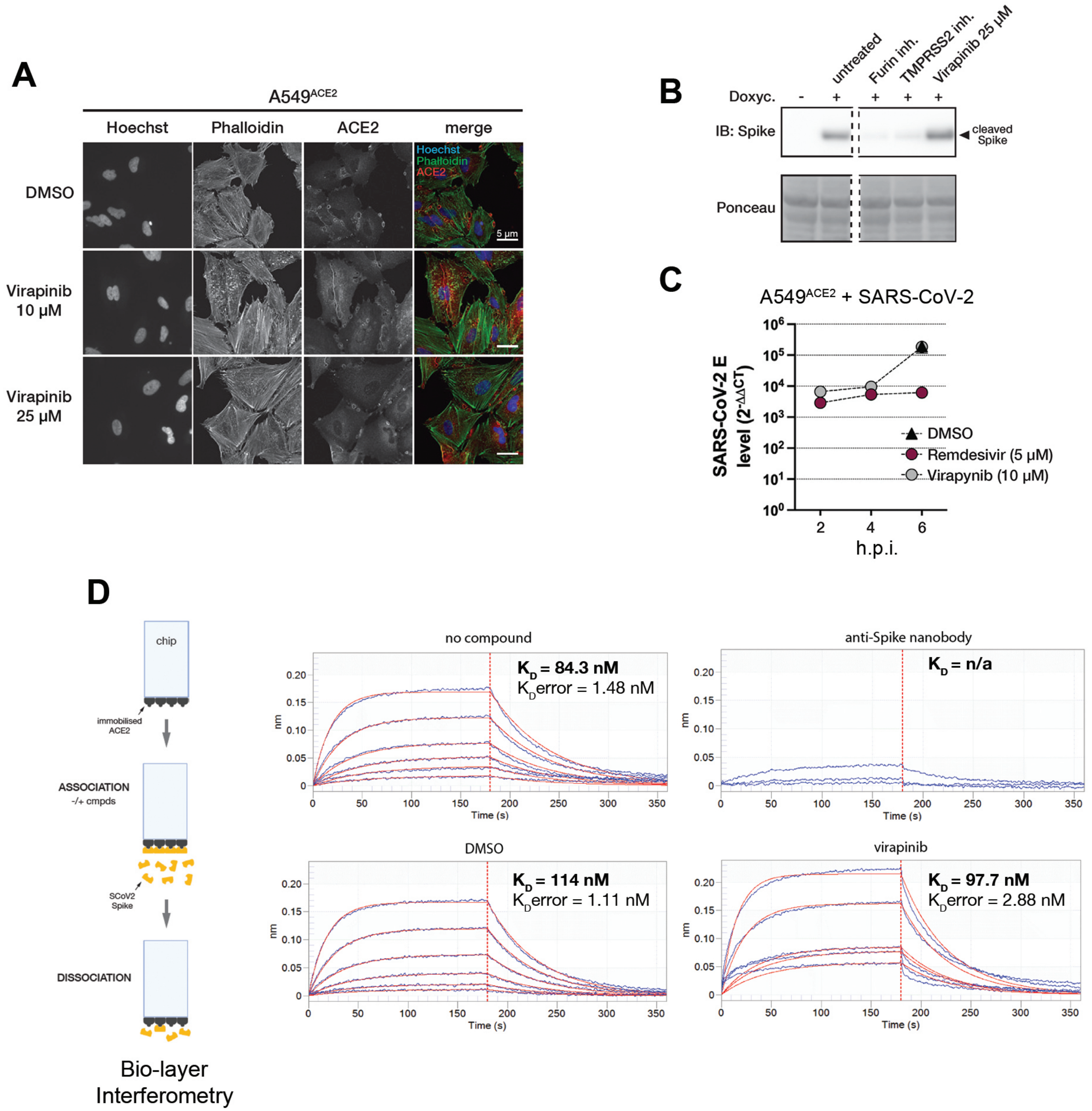
A
A549ACE2
Hoechst
Phalloidin
ACE2
merge
DMSO
Virapinib 10 µM
Virapinib 25 µM
5 µm
B
untreated
Furin inh.
TMPRSS2 inh.
Virapinib 25 µM
Doxyc.
- + + + +
IB: Spike
cleaved Spike
Ponceau
C
A549ACE2 + SARS-CoV-2
SARS-CoV-2 E level (2-ΔΔCT)
h.p.i.
DMSO
Remdesivir (5 µM)
Virapynib (10 µM)
D
chip
immobilised ACE2
ASSOCIATION
-/+ cmpds
SCoV2 Spike
DISSOCIATION
Bio-layer Interferometry
no compound
KD = 84.3 nM
KDerror = 1.48 nM
anti-Spike nanobody
KD = n/a
DMSO
KD = 114 nM
KDerror = 1.11 nM
virapinib
KD = 97.7 nM
KDerror = 2.88 nM
nm
Time (s)

**Figure S5. Effects of virapinib on ACE2, S protein and ACE2-S interaction.** (**A**) Representative images of ACE2 distribution. A549$^{ACE2}$ cells were treated for 24 h with indicated compounds, fixed and stained for ACE2 (red), Hoechst 33342 (blue) and Phalloidin (green). Scale bar (white) represents 5 μm. (**B**) WB analysis of the proteolytic processing of S protein. HEK 293T cells with dox-inducible expression of S protein were exposed to the indicated compounds for 6 h prior to a 16 h treatment with dox. Note that in contrast to Furin or TMPRSS2 inhibitors, virapinib does not affect the processing of the S-protein. Dashed lines indicate irrelevant lanes that have been omitted for clarity. The image of the Ponceau gel is shown to illustrate equivalent loading controls. (**C**) Binding curves from biolayer interferometry show association and dissociation of S to ACE2, which is bound to the sensor. A range of concentrations of S was used in each experiment, and compounds were added to test their effect on the ACE2-S interaction. In contrast to an anti-Spike nanobody control, virapinib does not impact the interaction kinetics or binding affinity. Binding curves (blue) show association signal (nm) on the y-axis and time (s) on the x-axis. Best-fit curves from a 1:1 binding model are shown (red). Note that the data from the anti-Spike nanobody could not be fit reliably, due to the dampening of the ACE2-S interaction, leading to low signal to noise ratios. (**D**) Analysis of SARS-CoV-2 replication. Cells were infected with SARS-CoV-2 for 1 h, after which the virus was washed off and compounds were added to the cells. Cells were collected for RNA extraction at 2, 4 and 6 h after infection. Viral RNA content was analysed by RT-qPCR. Plotted data represent mean the mean ± S.D. of 3 independent samples.
